# Rapid Derivation of Cloning-Competent Cells from Peripheral Blood Advances Conservation Biobanking

**DOI:** 10.1101/2025.10.08.680978

**Authors:** Laura Knecht, Anthony Mastracci, Tyler Miyawaki, Fatima Charles, Gregory L. Gedman, Riley Harbert, Jason Herrick, Laura L. Kuperman-Yao, Linda Lay, Alba Ledesma, Steve Metzler, Kathleen Morrill Pirovich, Dennis Milutinovitch, James B. Papizan, Parker Pennington, Lexie Russell, Kerry Ryan, Kanokwan Srirattana, Kaitlin Steiger, Darya Tourzani, Kelcey Walker, Rafael V. Sampaio, Sven Bocklandt, Ben Lamm, Matt James, Shawn Walker, Beth Shapiro, James Kehler

## Abstract

Establishing viable cell lines from endangered species is essential for conservation, yet traditional fibroblast derivation from skin biopsies faces challenges including variable success rates, contamination risk, and extended culture timelines. We demonstrate that endothelial progenitor cells (EPCs) and pericytes isolated from peripheral blood represent superior alternatives for biobanking across three mammalian genera (*Canis*, *Bison*, and *Equus*). Blood-derived cells exhibited 2-3 fold faster doubling rates (15-20 hours versus >35 hours for fibroblasts) and reduced time to generate banked lines from 3-4 weeks to 1.5-2 weeks. Proteomic profiling of 32 canonical markers confirmed EPCs and pericytes represent distinct populations with lineage-specific molecular signatures. Optical genome mapping demonstrated equivalent genomic stability across all cell types with no detectable structural variants or aneuploidies. Critically, interspecific somatic cell nuclear transfer (iSCNT) experiments confirmed both EPCs and pericytes generate viable embryos with efficiency meeting or exceeding fibroblasts. Gray wolf blood-derived cells produced six viable fetuses with 15% implantation rate, while bison EPCs showed higher blastocyst formation (7%) than fibroblasts (3%) from the same individual. Blood collection during routine veterinary procedures offers minimally invasive sampling with reduced contamination compared to skin biopsies. These findings support integrating blood-derived cell banking into conservation programs, enabling opportunistic genetic preservation during standard management activities and expanding options for genetic rescue through assisted reproductive technologies.

## II. Introduction

Primary cell derivation offers significant potential for species conservation by proactively safeguarding genetic diversity, a crucial resource for species facing population fragmentation, inbreeding, and extinction. Major biobanks, such as the Frozen Zoo^®^ at the San Diego Zoo Wildlife Alliance, have amassed substantial collections (Mooney et al., 2023). The Frozen Zoo, for example, holds over 10,000 individual cell lines representing more than 1,100 species and subspecies, including approximately 5% of threatened mammals, birds, amphibians, and reptiles listed on the IUCN Red List (https://sandiegozoowildlifealliance.org/pressroom/news-releases/san-diego-zoo-globals-frozen-zoor-hits-milestone-10000-cell-lines). Cryopreserved cells from endangered wildlife can be reprogrammed into stem cells (Hutchinson et al., 2024) and/or used for reproductive cloning by conspecific or Interspecific Somatic Cell Nuclear Transfer (iSCNT) into related domestic oocytes and surrogate species (Cowl et al., 2024; Fritts, 2022). This conservation cloning approach has been attempted in a wide range of amphibian and mammalian species (Novak, Brand, et al., 2025). Recent successful examples to expand the genetic diversity of captive breeding programs include the introduction of an unrepresented founder of Black Footed Ferrets, *Mustela nigripes* (Novak et al., 2024; Wisely et al., 2015), and the use of an historically under-represented male line of Przewalski’s Horse, *Equus przewalskii (Novak, Ryder, et al., 2025)*. Concerns over introduction of mitochondria from the domestic oocyte donors can readily be mitigated by only cloning male donors or crossbreeding for one generation, and only propagating F1 paternal lines (Cowl et al., 2024). The safeguard of controlled breeding can readily be incorporated to support the universal goals of maintaining and expanding the genetic diversity of populations within individual species survival plans and international protections and conservation planning promoted by the IUCN (https://www.cpsg.org/our-approach/one-plan-approach-conservation).

Tissue sampling techniques such as taking skin punch biopsies of restrained animals or using dart biopsies of free-ranging wildlife are two approaches to establishing fibroblast cultures (Plikus et al., 2021). However, the success of fibroblast derivation can vary dramatically across species. While some, like the Chilean shrew, have reported high success rates (e.g., 90% with dermatome punches; Tovar et al., 2008), others, such as elephants, exhibit low and inconsistent success (0%-80%) even from biopsies of the same individual (Jansen van Vuuren et al., 2023). In addition, skin samples are covered with commensal microbes that can contaminate cell cultures, despite efforts to reduce this risk in the clinic, field and lab.

In an effort to establish a less invasive sampling technique for bio-banking, genomic, and iSCNT studies, we isolated, expanded, and characterized adherent stable cell populations from peripheral blood samples from several species. Endothelial progenitor cells (EPCs) are a rare (∼1 in a million) cell type circulating in peripheral blood that have been previously isolated and cultured from blood from mice to humans (Jansen van Vuuren et al., 2023; see review by Yoder, 2012). Pericytes are believed to reside on the outside of the endothelial layer of small blood vessels such as capillaries, interacting with EPCs (Alarcon-Martinez et al., 2021; Attwell et al., 2016) and are typically isolated from dissected and digested tissues (McErlain et al., 2025). Here, we were surprised to discover that we could expand both EPCs and pericytes from peripheral blood samples using the same method. It is possible that prior cultures of presumptive EPCs may have been misidentified or also contain pericytes. Despite this potential admixture, we demonstrate that cultures of both non-hematopoietic cell populations from blood can be isolated and expanded for biobanking and cloning in conservation.

## III. Methods

### A. Blood Derivation and Culture

We collected blood samples from four species: gray wolf (*Canis lupus*) and domestic dog (*C. l. familiaris*), bison (*Bison bison*), horse (*Equus caballus*), and all major zebra species (*Equus quagga*, *E. zebra*, and *E. grevyi*). Hereafter, we refer to these as canids, bovids, and equids. All samples were collected in ACD blood tubes and transported overnight at room temperature.

After recording blood volume, we diluted each sample with 1xdPBS (Cat#: 14190-144;Fisher Scientific) and 3% HiFBS (Cat#: 16140071; Gibco) at a 1:3 ratio. We then gently and equally layered the diluted blood on top of 15mL of Ficoll Paque Plus (Cat#: 95021-205; VWR) in 50mL conical tubes. For example, 60mL of diluted blood would be split into two 50mL conicals with 15mL Ficoll Paque Plus per conical. We then centrifuged the dilute blood:Ficoll Paque Plus at 800g for 25 minutes at room temperature with the break set to 0, and then aspirated the plasma layer. We then transferred the PBMC layer to another 50mL conical tube and brought the total volume up to 50mL with dPBS. We then centrifuged the conical tube at 800g for seven minutes with the brake set to 9. We aspirated the supernatant and then resuspended it in 30mL of dPBS, and centrifuged at 400g for 5 minutes. We then aspirated the supernatant and resuspended it in 1mL of EGM (20% HiFBS, + EGM (Cat#: CC-3124; Lonza). The cell suspension was then counted using AOPI Cell Staining Solution (Cat#: CS2-0106-25mL; Fisher Scientific), and resuspended in 25mL of EGM and transferred to a Biocoat Collage T175 Flask (Cat#: 356487; Corning) and incubated at 37C. The next day, we changed the media, and then continued media changes every other day until colonies had formed. Once colonies appeared, we performed daily half media changes until the colonies were 5mm in diameter or were greater than 20 total colonies per flask.

We lifted the plate using 50:50 ratio of 0.05% Trypsin (Cat#: 25300-054; Fisher Scientific) and Accutase (Cat#: SF006; Millipore) for 3-5 minutes at 37C. Once cells were lifted, we quenched the solution with EGM and then centrifuged it at 400g for 5 minutes. The media was aspirated and then resuspended in 1mL of EGM. We then counted cells using AOPI and plated them onto Biocoat Collagen T175 Flasks and seeded at 1-2x10^6 per flask. When the cells were confluent, we repeated the same procedure and, instead of the cells being replated, they were centrifuged at 400g for 5 minutes and then resuspended at a concentration of 1x10^6/mL in EGM freezing media (30% EGM, 60% HiFBS, and 10% DMSO (Cat#: D2650-5x5mL).

### B. Fibroblast Derivation and Culture

We obtained dermal tissue from ear notches for fibroblast cell line derivation. To account for contamination from the field, we divided the samples in half and generated an explant derived cell line and a digested derived fibroblast cell line. We generated digested derived fibroblasts by mincing in collagenase IV (Cat#: C4-28-100MG; Sigma) and then incubating at 37C for 1hr. The cells were then washed and plated onto a tissue culture treated T75 flask (Cat#: 229340; CellTreat) in cDMEM [DMEM high glucose, + sodium pyruvate, - glutamine (Cat#: 10313-021; Fisher Scientific), 10% FBS, 1X Glutamax (Cat#: AI2860-01; Fisher Scientific), 1X MEM-NEAA (Cat#: 11140-050; Fisher Scientific), 1X 2-mercaptoethanol (Cat#: 21985023; Gibco)]. The cells were then cultured, passaged, and frozen at p2 to p3 in FB Freezing Media (30% cDMEM, 60% HiFBS, and 10% DMSO).

### C. Doubling Rate

We used the following format to calculate the doubling rate across all species and cell types: doubling rate= hours in culture * LN(2)/LN(final cell number/initial cell number).

### D. Proteomics

We maintained cells under standard culture conditions until passage 5 and cultured to 80% confluency in two six-well plates for biological replicates. On the day of lysis, we gently washed cells twice with 1 mL of 1× dPBS, and discarded the supernatant. We lysed the cells by adding 75 µL of RIPA buffer (Cat# J63306.AP) pre-mixed with HALT protease inhibitor (Cat# 87785). Subsequently, we incorporated 1 µL of nuclease (Cat# 88700) and mixed post-lysis. We quantified total protein concentration using Thermo Fisher’s BCA Gold assay (Pub. No. MAN0029413 Rev. B.0). Lysates were stored under -80C in 1.5mL low bind Eppendorf tubes until all samples were collected for downstream mass spectroscopy library preparation.

We generated peptides via trypsin digestion and labeled using the TMT10plex isobaric labeling kit (Cat# 90406; Thermo Fisher). We pooled labeled samples by species and subjected each pool to high-pH fractionation to yield eight fractions. We analyzed each fraction using an Orbitrap Eclipse mass spectrometer processed with Proteome Discoverer 3.0 software. Peptides were annotated against species-specific UniProt databases. Following data acquisition, we filtered proteins based on false discovery rate (FDR), retaining only those with an FDR <1% and a normalized average intensity >2000 units across grouped samples for downstream analysis.

### E. Reference Genome Assemblies

We downloaded genome assemblies from GenBank for mapping. For canids, we used the gray wolf assembly GCA_905319855.2, which includes autosomes, sex chromosomes, and a mitochondrial genome. For equids, we used the domestic horse assembly GCA_041296265.1, which includes autosomes, chromosome X, and a mitochondrial genome. For bovids, we combined the autosomes, X chromosome, and mitochondrial genome from bison assembly GCA_030254855.1 (a female) and the bison Y chromosome from the F1 cattle x bison assembly GCA_018282365.1.

### F. FACs Staining and Analysis

We harvested 3x10^5^ cells per condition per line and then washed cells for 30 minutes with cold blocking buffer [90% dPBS (Cat#: 14190-144;Fisher Scientific) and 10% Goat serum (Cat#: 16210-072; VWR)]. The cells were then pelleted and washed twice with dPBS at 400g for 5 minutes. We incubated cells for 1 hour at room temperature in the dark with 200uL of Flow Cytometry Antibody Dilution Buffer (Cat#: 13616; Cell Signaling Technology) plus the corresponding antibody. We diluted the Rat IgG2b Isotype Control (APC) (Cat#: 17-4031-82; eBioscience) and Mouse IgG1 Isotype Control (PE)(Cat#: 12-4714-82; eBioscience) 1:200, and added the CD90 (APC) (Cat#: 17-5900-42; eBioScience) at 1:20, CD34 (PE) (Cat#: 12-0340-42; eBioscience) to make 3µg/mL. The dual stain condition was the same with CD90 at a 1:20 dilution and CD34 at 3µg/mL dilution. After incubation, we pelleted cells and washed three times with dPBS at 400g for 5 minutes. We resuspended cells into 200uL of cold FACs buffer (dPBS, 2% FBS(Cat#: 16140071; Gibco), 2.5mM EDTA pH 8.0(Cat#:15575020; Thermofisher), 25mM HEPES (Cat#: 15630-080; Fisher Scientific), and 1% Pen/Strep 100X (Cat#: 15140122; Fisher Scientific)and filtered into a 5mL FACs tube (Cat#: 196-19881-083; Spectrum Laboratory Products). We then analyzed cells on BD FACSDiscover™ S8 Cell Sorter, with a VBYR laser configuration, 100µm nozzle size, and standard configuration and settings while using the YG1 (575) and R1 (655) lasers.

### G. Angiogenesis

We thawed Matrigel (Cat#: 354230; Corning) on ice and then plated 75uL per well of a 96 well plate. The plate was then incubated at 4°C for at least 1 hour. We gently aspirated the Matrigel prior to plating (For blood derived cells: Cat#: 354409; Corning; For fibroblasts: Cat#229196; Cell Treat), and then incubated at 37°C for 10 minutes. Cells were lifted by normal culture conditions, and plated at 1x10^4 for EPCs and Pericytes, and 2x10^4 for fibroblasts in duplicates into both Matrigel coated wells and uncoated 96 wells with 100 µL media. The plate was then incubated at 37°C for a maximum of 24 hours, and imaged at least 4 times over the time period. In general, the assay showed tubule formation by 12 hours.

### H. Low Density Lipoprotein & Acylated Low Density Lipoprotein Uptake

We grew fibroblasts and blood derived cells under standard cell culture conditions until the cell lines had reached 65-80% confluency. On the day of passaging, we collected 1.0x10^5 cells from each cell line, pelleted and washed cells with 1X DPBS, and then resuspended them in serum free DMEM (DMEM, -FBS, 0.3% MACS® BSA (Cat#: 130-091-376, Miltenyi Biotec)) for fibroblasts and serum free EGM (EGM,-FBS, 0.3% MACS® BSA) for blood-derived cells. We then transferred cells onto a Corning 96-well for blood-derived cells and Cell Treat 96-well for fibroblasts with 1.0x10^4 cells per condition. The plates were then incubated at 37°C for 16 hours. We washed the cells with LDL Assay buffer (1X DPBS, +calcium, +magnesium (Cat#: 14040133; Gibco), 0.3% MACS® BSA) and resuspended in serum free DMEM for fibroblasts and serum free EGM for blood derived cell lines. Low Density Lipoprotein from Human Plasma, BODIPY™ FL complex (Cat#: L3483, Invitrogen™) was diluted 1:100 for a final concentration of 10 µL/mL and added to the appropriate wells in replicates. Low Density Lipoprotein from Human Plasma, Acetylated, Alexa Fluor™ 594 Conjugate (Cat#: L35353; Invitrogen) was also diluted 1:100 for a final concentration of 10 µL/mL and added to the appropriate wells in replicates. Cells were returned to the incubator at 37°C for 2.5 hours. After incubation, we washed the cells three times with the LDL Assay Buffer and then imaged and analyzed on the Incucyte® SX5 utilizing the Orange/Green optical modules.

### I. Karyotyping

We shipped live canid (gray wolf) cell cultures to an independent cytogenetics laboratory (KaryoLogic Inc.) in 2 T25 flasks at 25% and 50% confluency so that they would arrive subconfluent, thus minimizing contact inhibition of the cells. Mitotic spreads were prepared and G-banded karyotypes assembled from 20 cells per line to determine the average karyotype and detect the presence of any aneuploidies.

### J. Optical Genome Mapping

We performed Optical Genome Mapping (OGM) analyses of the derived cell lines using the Bionano Saphyr platform according to manufacturer specifications. Briefly,we used 500k-750k cells to isolate ultra-high molecular weight (UHMW) genomic DNA as described in the Bionano Prep SP-G2 Fresh Cell Pellet DNA Isolation Protocol (Bionano, Doc no. CG003). We then fluorescently labeled UHMW gDNA by the Direct Label Enzyme (DLE-1) and then stained for backbone visualization (Bionano Prep Direct Label and Stain (DLS) Protocol, Doc no. 30206). We loaded the labeled and stained UHMW gDNA sample onto a Saphyr Chip and into the Saphyr System Instrument (Bionano Saphyr System User Guide, Doc no. 30247). We used Bionano Solve software to identify structural variations (SV) and low variant allele fraction variants (VAF) (Bionano Solve Theory of Operation: Structural Variant Calling, Doc no. CG-30110). The resolution for detecting variants with this method was >500bp up to the chromosome level, when confirmed by short-read WGS assemblies.

### K. Interspecies Somatic Cell Nuclear Transfer

Synchronization of fibroblasts, EPCs and pericyte donor cells for iSCNT was achieved by allowing cultures to grow to complete confluence to induce contact inhibition and enter cell cycle (G0) arrest. For gray wolf iSCNT, we used either EPCs or pericytes from *Canis lupus* as nuclear donors. We isolated oocytes from donor dogs in estrus and performed enucleation, reconstruction, electroporation, subsequent activation, and overnight *in vitro* culture (IVC) as described previously (G. A. Kim et al., 2012). For bison we isolated cattle cumulus oocyte complexes (COCs) from abattoir ovaries for subsequent *in vitro* maturation (IVM), enucleation, reconstruction with female bison EPCs or fibroblasts and activation as previously published in detail for cow SCNT (Sangalli et al., 2023). Reconstructed Bison iSCNT embryos were then cultured in a bovine IVC medium for 7 days and cryopreserved at the blastocyst stage (Barfield, 2019; Benham et al., 2021). For equids, individual donor EPCs and fibroblasts from a male plains zebra were injected into the ooplasm of enucleated domestic horse oocytes obtained by ovum pick-up, electrofused, and activated to produce reconstructed iSCNT plains zebra embryos as performed in the successful generation of cloned Przewalski’s Horse by iSCNT (Novak, Ryder, et al., 2025). Reconstructed plains zebra embryos were cultured and frozen at the morulae and blastocyst stages 6-8 days post cloning (Hinrichs, 2020).

## IV. Results

### A. Derivation Success and Timeline

We successfully derived adherent cell populations from peripheral blood samples across three genera: *Canis,* including gray wolf and domestic dog, *Bison*, where all individuals were plains bison, and *Equus*, which included domestic horse and major zebra species (Table 1). We found that consistent successful derivation of canid cells required a minimum of 15mL of blood, while bovid and equid cell lines required at least 25mL of blood. Blood-derived cell cultures showed an overall derivation success rate of 78% (Table 1). Time to initial colony formation varied by species (Fig. 1): bison formed colonies in 4±1 days, equids in 6±1 days, and gray wolf in 7±1 days post-processing. Due to rapid proliferation, all blood-derived cultures generated 30 million low-passage (P1-2) cells within 1.5-2 weeks of initial plating.

**Figure 1.**
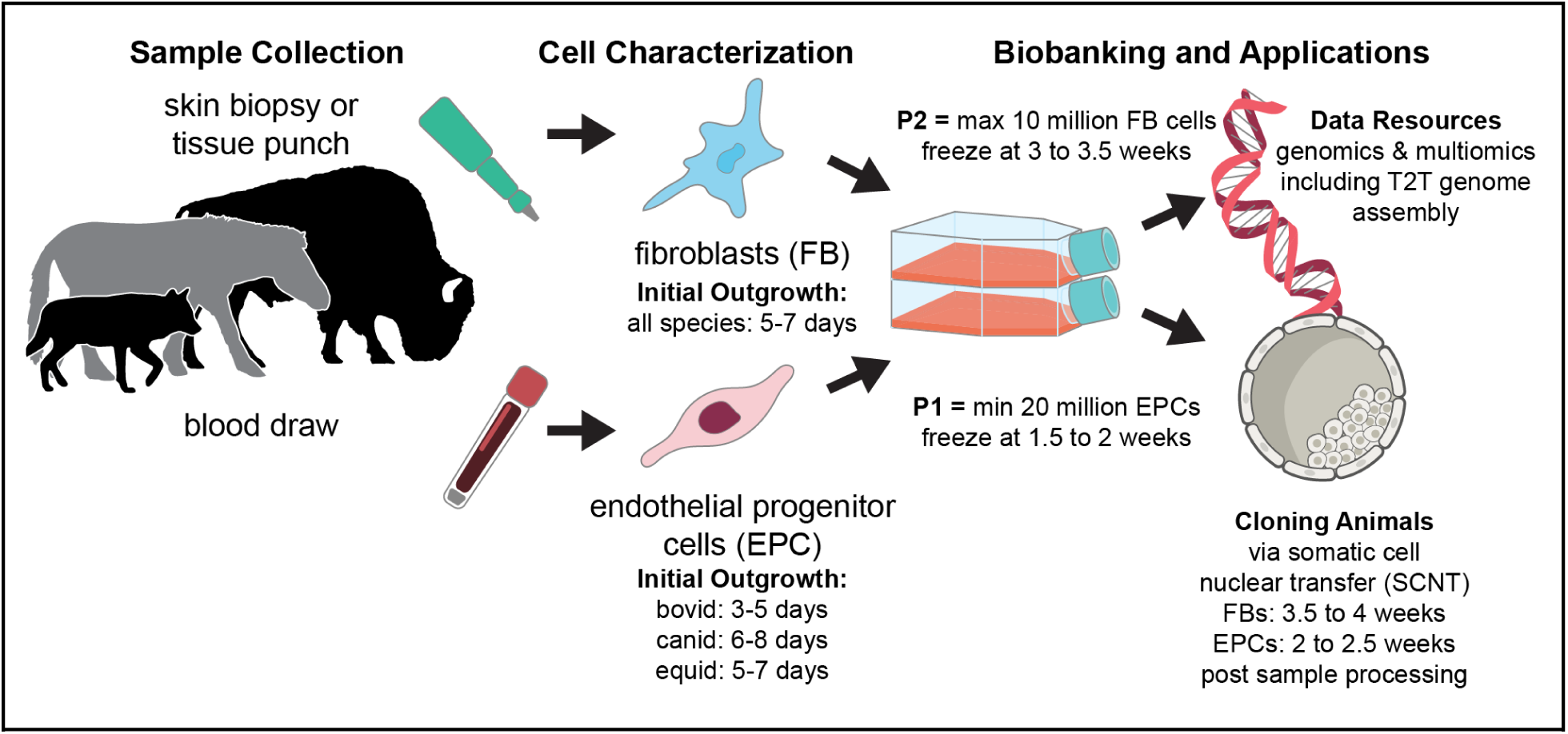
Cell lines can be generated from both fibroblasts and cells (pericytes and endothelial progenitor cells, or EPCs) isolated from blood draws. Cells isolated from blood show an expedited timeline to handoff for downstream applications compared to fibroblasts.

**Table 1.** Cell line derivation success rates by species and source.

| Genus species | Blood-derived culture |  |  | Ear Notch Fibroblast culture |  |  |
| --- | --- | --- | --- | --- | --- | --- |
|  | Success | Failure | % Success | Success | Failure | % Success |
| *Canis lupus* | 2 | 0 | 100% | 3 | 0 | 100% |
| *Canis lupus familiaris* * | 2 | 1 | 67% | 0 | 0 |  |
| *Bison bison* | 12 | 3 | 80% | 13 | 4** | 76% |
| *Equus caballus* | 3 | 1 | 75% | 0 | 0 |  |
| *Equus grevy* * | 0 | 1 | 0% | 1 | 0 | 100% |
| *Equus zebra hartmannæ* | 1 | 0 | 100% | 1 | 0 | 100% |
| *Equus quagga* | 1 | 0 | 100% | 1 | 0 | 100% |
| Total Success Rate | 21 | 6.00 | 78% | 19 | 0 | 95% |
| *Excluded fetal skin |  |  |  |  |  |  |
| **Failed due to contamination |  |  |  |  |  |  |

### B. Growth Characteristics, Morphology, and Surface Markers

Doubling rates differed consistently between cell types across species (Fig. 2). EPCs exhibited the fastest average doubling rate at 15 hours, pericytes averaged 20 hours, and fibroblasts showed more variable rates exceeding 35 hours. Morphologically, fibroblasts presented as elongated spindle-shaped cells with central nuclei across all genera (Fig 3). EPCs displayed small, cobblestone-like morphology with high cell aggregation. Pericytes showed fibroblast-like morphology but with less cellular elongation (Fig 3).

**Figure 2.**
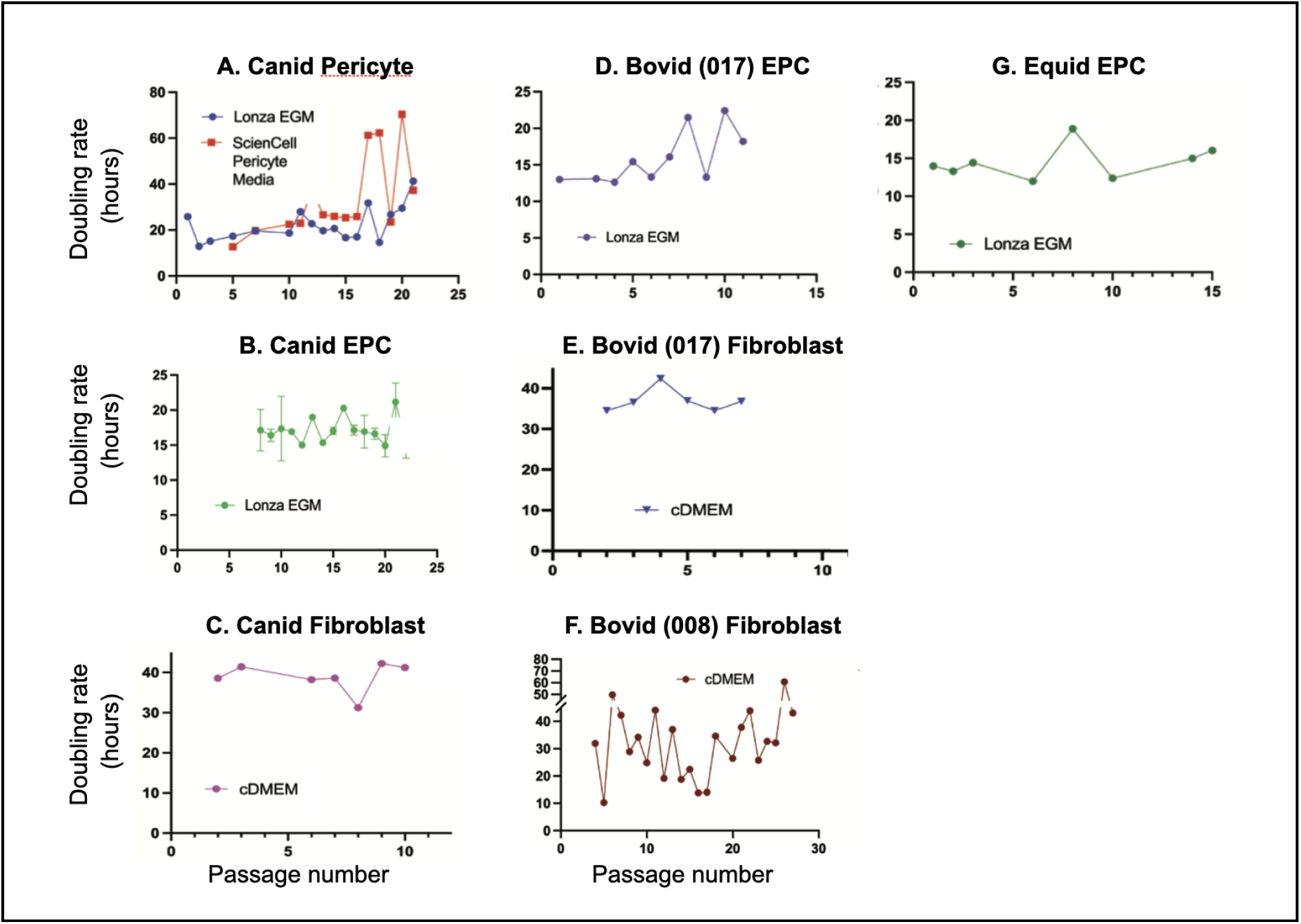
Doubling times by passage for canid **(A-C)**, bovid **(D-F)**, and equid **(G)** cells. Data was collected via Incucyte Cell imaging with the Cell-by-Cell AI Analysis software trained to analyze each cell type. Cells were imaged daily to capture live cell count, which was used to calculate the average doubling rate across each passage.

**Figure 3.**
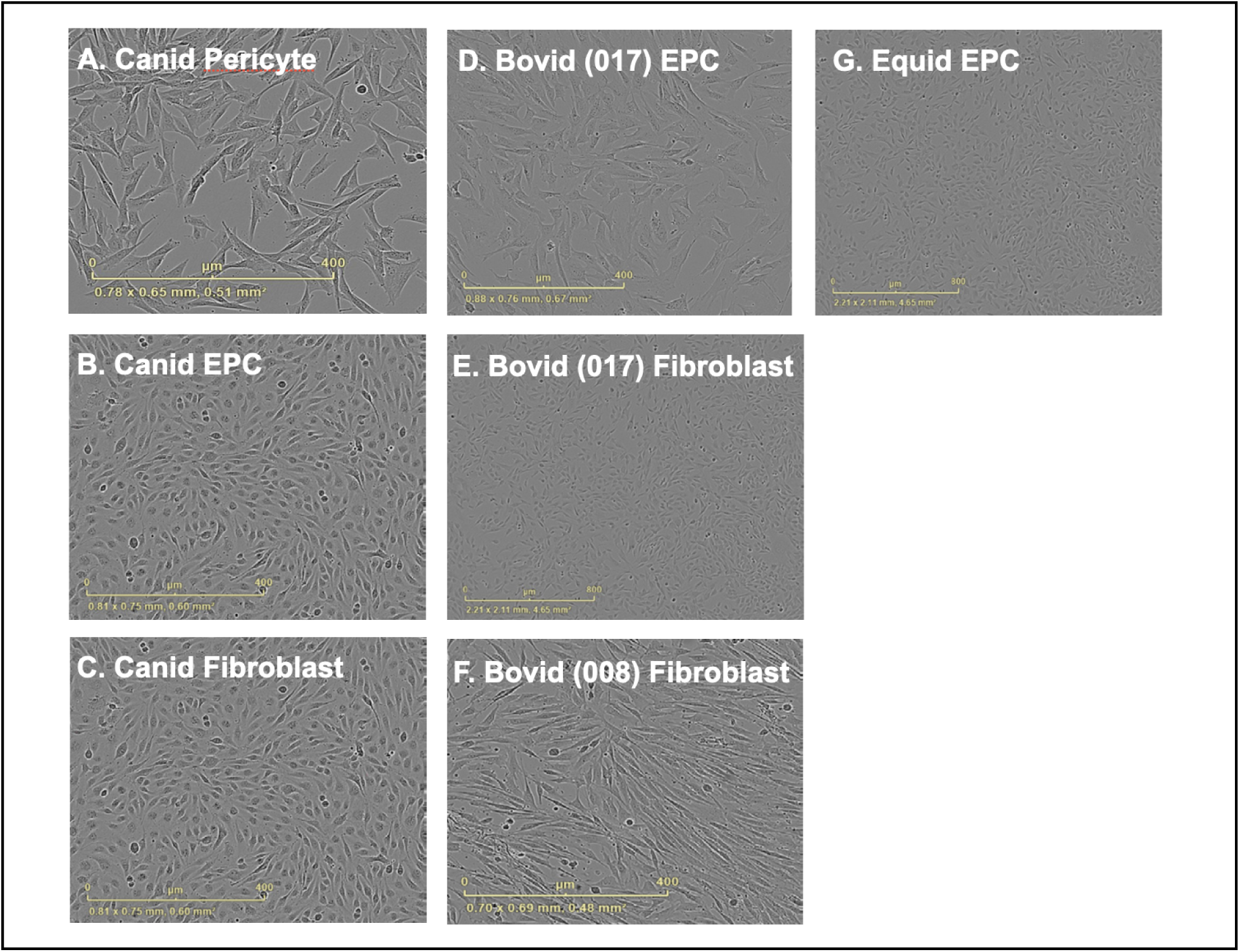
Morphology of canid **(A-C)**, bovid **(D-F)**, and equid **(G)** cells. All photos were taken under the phase imaging of the Incucyte. Images A-D & F were captured with the 10X objective, while images E and G were captured with the 4X objective.

Flow cytometry analysis of canid cells using anti-dog CD34 and CD90 antibodies showed distinct surface marker profiles (Fig. S1). EPCs contained a discrete 25% population of CD34+ cells. Pericytes contained a minor ∼12% sub-population of CD90+ cells with negligible (< 1%) detection of CD34. Fibroblasts were uniformly 100% CD90+ with a minor 3.5% sub-population % CD34+ cells, possibly reflecting heterogeneity in early passage cultures.

### C. Functional Characterization

In vitro angiogenesis assays demonstrated that EPCs formed tubule-like networks when plated on extracellular matrix substrate, while fibroblasts showed no morphological changes (Fig. 4A). Pericytes adhered to the outer surface of EPC-derived tubule structures in co-culture experiments. Neither pericytes nor fibroblasts formed tubules when plated alone.

**Figure 4.**
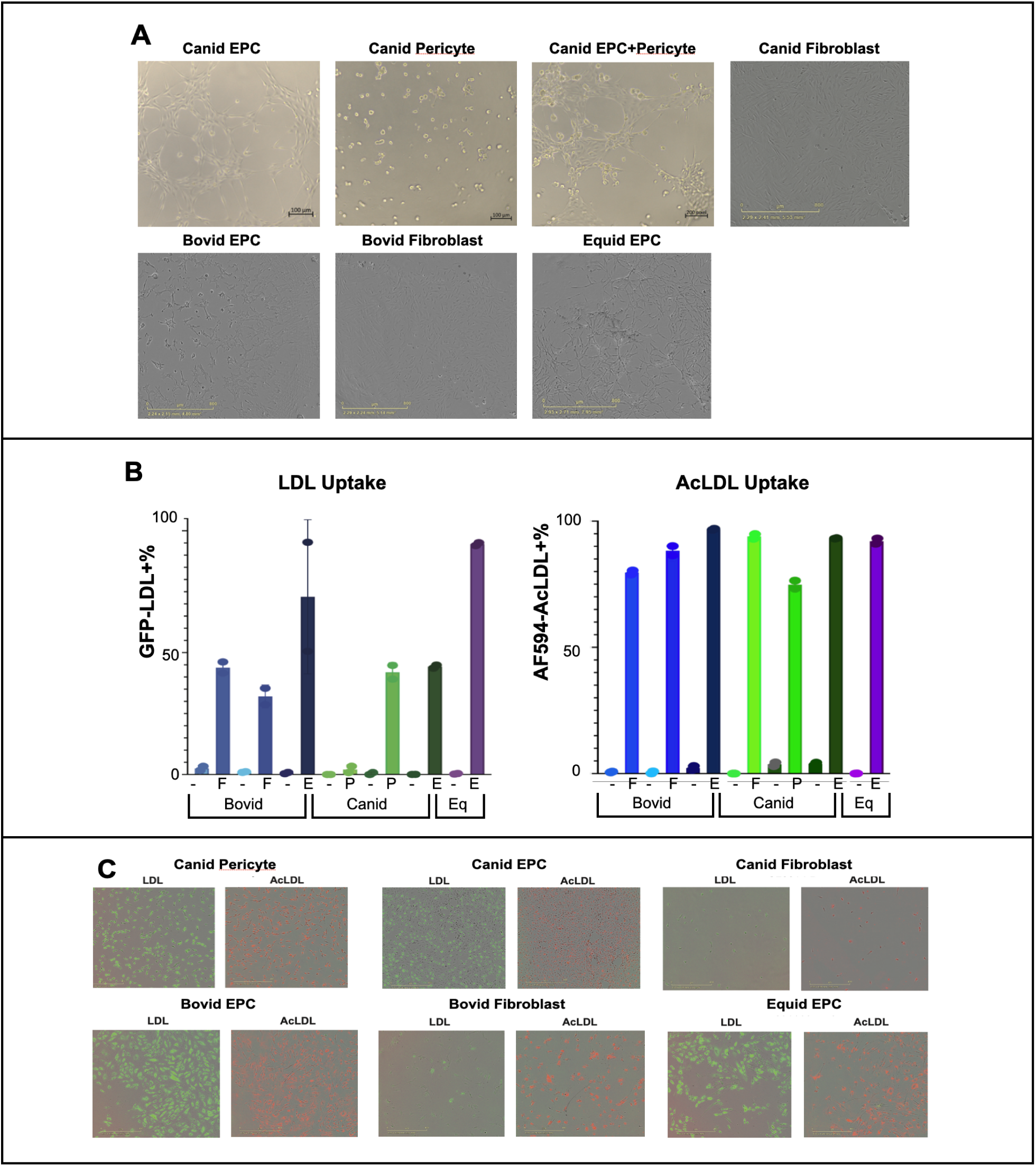
Functional assays for cell characterization. (**A**) Results of the angiogenesis assay. Canid EPC, pericyte, and EPC+pericyte images were taken using the Zeiss Primovert with a 10x Objective. The remaining images were taken using the Incucyte at a 10X objective. EPCs formed tubule networks, while pericytes adhered to the outer layer of the EPC-tubule network. Pericytes and fibroblasts plated alone did not form tubules. (**B**) Results of the Low Density Lipoprotein & Acylated Low Density Lipoprotein Uptake assay. A “-” below a bar indicates negative controls, while E, P, and F indicate EPCs, pericytes, and fibroblasts respectively. Cell fluorescence was measured using Incucyte Cell-by-Cell AI Analysis software trained on these cell types. (**C**) Fluorescent microscopy images of cells taken by the Incucyte at a 10x objective with the Green and Orange fluorescence channels to generate positive cell fluorescence data. Fibroblasts have a lower level of BodipyFL-tagged (Green) LDL uptake than those of blood derived cell lines, while pericytes have lower levels of AlexaFluor594-tagged (Red) AcLDL uptake than EPCs and fibroblasts from the same species.

Low Density Lipoprotein (LDL) and Acetylated-LDL (AcLDL) uptake assays revealed differential internalization patterns (Fig. 4B,C). Both EPCs and pericytes showed elevated LDL uptake compared to fibroblasts. For AcLDL uptake, EPCs demonstrated highest uptake, followed by fibroblasts, with pericytes showing the lowest uptake levels. This differential AcLDL uptake can be used to distinguish EPCs from pericytes in cultures originating from blood samples.

### D. Genomic Stability Assessment

Optical genome mapping (OGM) with ∼80-fold genomic coverage confirmed expected chromosome copy numbers and sex chromosome compositions for all cell lines (Fig. 5, Fig. S2) OGM data from male canid samples was concordant with independent G-banded karyotype analysis (Fig. 5). Both male gray wolves from which cell lines were derived had 38 pairs of autosomes and one X and one Y chromosome. Bovid samples had 29 pairs of autosomes and X chromosome copy number consistent with known sex (Female 008; Male 017 EPC and fibroblasts in Fig. S2A). Equid samples had 31 pairs of autosomes and a pair of X chromosomes, consistent with female sex (Fig. S2B). No significant structural variants or aneuploidies were detected, confirming karyotypically normal primary cells suitable for downstream applications.

**Figure 5.**
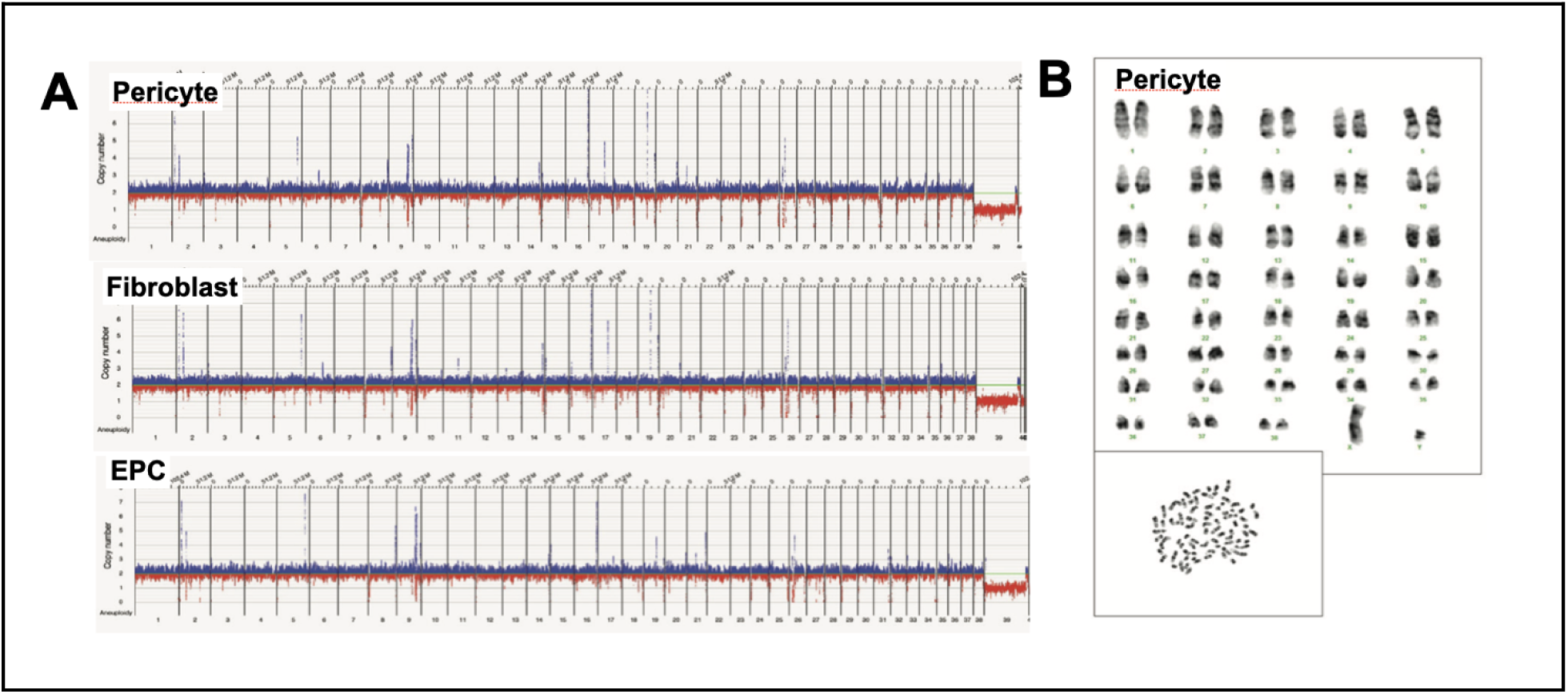
A. Optical Genome Mapping (OGM) plots summarizing chromosome copy numbers for canid cell lines. **B.** G-banded karyotype analysis of canid pericytes showing normal karyotype.

### E. Proteomic Characterization of Cell Populations

Analyses of gene expression data from 32 canonical cell markers revealed distinct clustering of the three primary cell populations: endothelial progenitor cells (EPCs), fibroblasts (FBs), and pericytes (Fig. 6). We used canid cell lines as a framework for our analyses, as canid annotations in the UniProt database both captured more genes and and higher annotation scores compared to other taxa. We performed species-specific analyses for canid and bovid samples only, due to the absence of comparative data from fibroblasts for horse or zebra.

**Figure 6.**
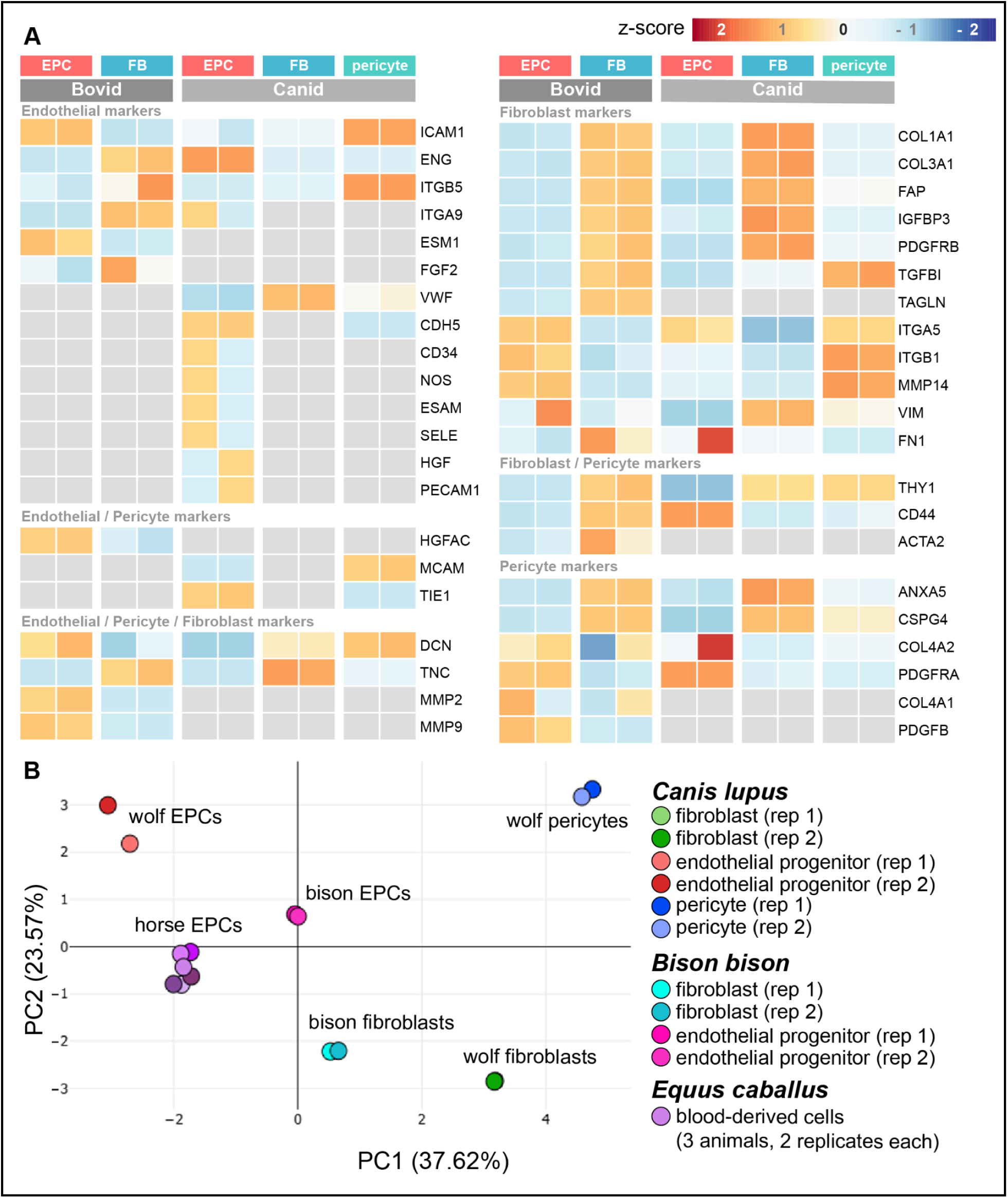
Results of proteomic analyses for replicate samples of each cell type. **(A)** Heatmap showing relative expression of canonical protein markers for cell type determination in canid and bovid cells. Protein expression was normalized, and color intensity reflects z-scores from -2.58 (blue, lowest) to 2.58 (red, highest); gray indicates undetected protein. Log2(fold-change) and -Log10(p-value) for each gene are provided in Table S1; **(B)** PCA of all samples of all three taxa using 18 canonical markers present in all cell lines: VIM, COL1A1, ANXA5, COL3A1, CSPG4, ITGA5, ITGB1, CD44, ENG, ICAM1, PDGFRA, MMP14, THY1, COL4A2, PDGFRB, DCN, TGFBI, and ITGB5. For equine samples, only blood-derived cells were available. Although the first two principal components account for 61% of the total variance, the PCA shows equine samples aligning more closely with canid and bovid EPCs than with their respective fibroblast or pericyte populations.

*Fibroblasts*. Canid and bovid fibroblasts exhibited elevated expression of extracellular matrix (ECM) proteins and markers associated with activated fibroblasts, including Fibroblast Activation Protein alpha (FAP), which is an established marker for activated fibroblasts in tissue remodeling (Tillmanns et al., 2015), and Insulin-like growth factor-binding protein 3 (IGFBP3), which is involved in modulating IGF signaling and ECM interactions. We observed a high abundance of major structural ECM components Collagen Type I Alpha 1 Chain (COL1A1), Collagen Type III Alpha 1 Chain (COL3A1), which aligns with the known matrix-synthesizing capacity of FBs. Vimentin (VIM), which is crucial for cell structure, and Platelet-Derived Growth Factor Receptor Beta (PDGFRB), a key receptor in fibroblast proliferation, as well as Tenascin C (TNC), an ECM glycoprotein induced during tissue injury and remodeling (Midwood et al., 2011). When comparing EPCs to FBs, Log₂(Fold-Change) values for these markers (Table S1) were consistently negative and statistically significant (Log₁₀(p-value) > 2.8), with more pronounced differences observed in canid samples (e.g., COL1A1: -6.2) compared to bovid samples (COL1A1: -1.1).

*Pericytes.* Pericytes demonstrated a high abundance of proteins involved in cell adhesion and ECM interaction, including Integrin beta 5 (ITGB5) and Intercellular adhesion molecule 1 (ICAM1), both of which are critical for cell-cell and cell-matrix interactions (Pang et al., 2018). High expression of Matrix Metalloproteinase-14 (MMP14) suggested potential for ECM degradation and turnover, key functions in vascular remodeling. Other highly expressed proteins include Transforming Growth Factor Beta Induced Protein (TGFBI) and Decorin (DCN), suggesting active ECM engagement,and Melanoma Cell Adhesion Molecule (MCAM/CD146), an important surface molecule on pericytes that downregulates TGFB signaling to impact endothelial cells junction integrity (Chen et al., 2017). In addition to fibroblasts, PDGFRB is a canonical marker of human and mouse pericytes involved in their recruitment and activation (reviewed by Yao, 2022). PDGFRB was abundant in canid pericytes and negligible in EPC.

*EPCs.* EPCs exhibited markers consistent with endothelial-like cells, including high expression of Endoglin (ENG), a TGF-β co-receptor, in both canid and bovid samples, and Cadherin 5 (CDH5/VE-Cadherin), which is essential for endothelial cell-cell junctions (Siavashi et al., 2016), detected specifically in canid EPCs. Platelet-Derived Growth Factor Receptor Alpha (PDGFRA) is expressed during fetal cardiac development and in rare post-natal cardiac endothelial cells (Chong et al., 2013) Interestingly, EPCs from both canid and bovid samples had abundant expression of PDGFRA when compared to species-specific fibroblasts. Canid EPCs expressed canonical EPC markers (reviewed by Hassanpour et al., 2023) including CD34, PECAM1, SELE, NOS3, ITGA9, ESAM, HGF, and ICAM1 in abundance that were not detected in canid fibroblasts or pericytes (Caiado & Dias, 2012; Liu et al., 2019). However, many canid EPC markers lacked identifiable gene orthologs that passed mass spectrometry filtering in the UniProt databases for bison or horses, limiting the assessment of their expression conservation for these species.

A cross-genera PCA including equid samples used data from 18 markers quantified across all cell lines (VIM, COL1A1, ANXA5, COL3A1, CSPG4, ITGA5, ITGB1, CD44, ENG, ICAM1, PDGFRA, MMP14, THY1, COL4A2, PDGFRB, DCN, TGFBI, ITGB5; the other 14 were excluded due to missing data; Fig. 6B). Equine blood-derived cells clustered more closely with canid and bovid EPCs than with fibroblast or pericyte populations. Some marker expression patterns deviated from canonical assignments (e.g., low VWF in EPCs, high VWF in canid fibroblasts; variable FN1 and CD44 expression across species and cell types), reflecting context-dependent expression profiles.

### F. Interspecies Somatic Cell Nuclear Transfer (iSCNT)

*Canids.* Both pericytes and EPCs successfully generated viable iSCNT gray wolf embryos. Twenty-one embryos reconstructed from pericytes and 18 from EPCs were transferred into a single recipient domestic dog in estrus (Fig. 7A,B). Transabdominal ultrasound at Days 21 and 23 post-embryo transfer confirmed implantation of six viable embryos with active heartbeats (Fig. 7G). Timed ovariohysterectomy at Day 33 yielded six morphologically normal fetuses. Genotyping using a custom SNP panel identified four fetuses derived from pericytes and two from EPCs, demonstrating viable implantation rates of 19% (pericytes) and 11% (EPCs).

**Figure 7.**
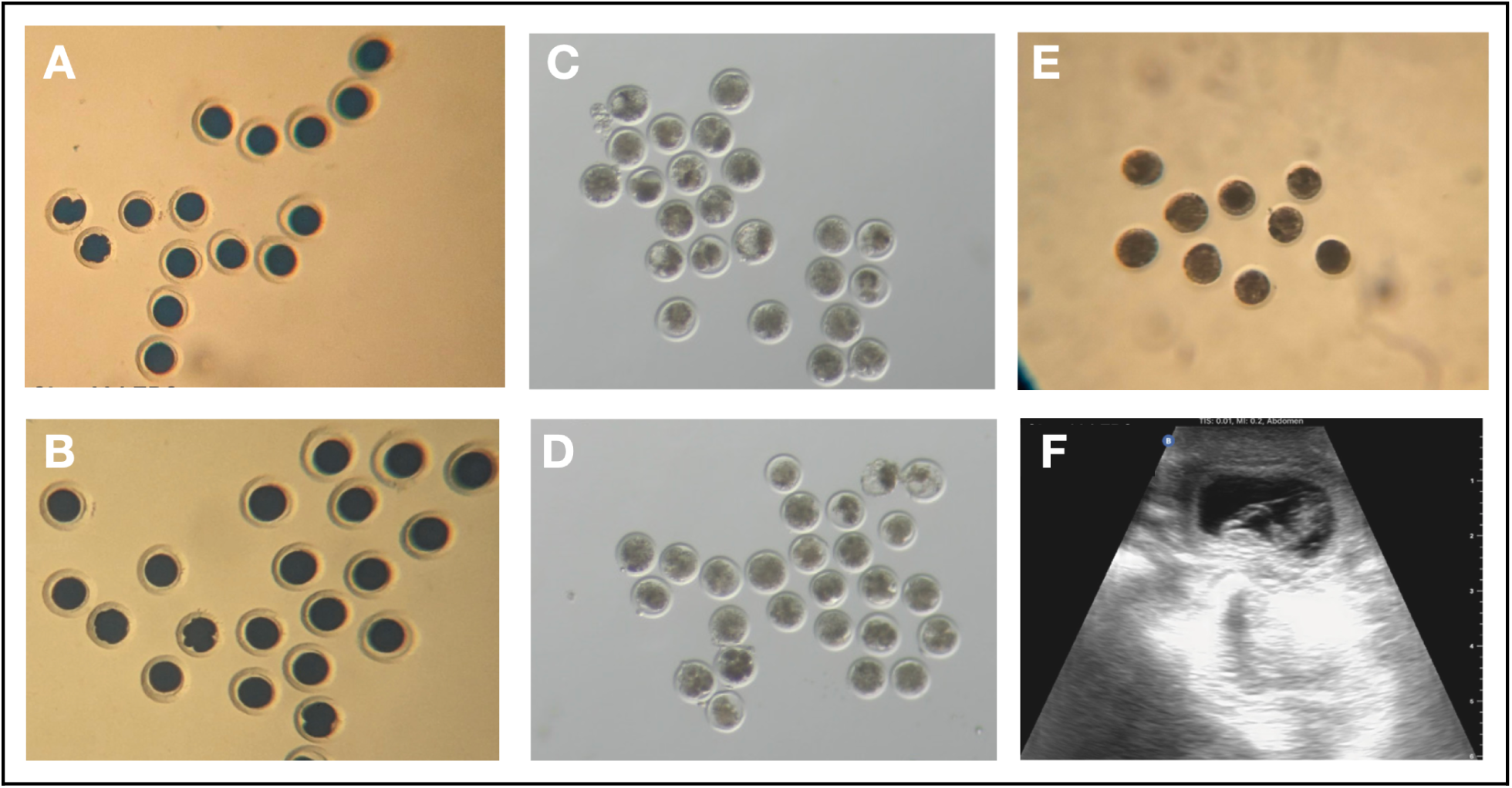
Brightfield images of pre-implantation embryos and post implantation ultrasound images. **(A)** Early cleavage stage embryos are visible in overnight cultures of iSCNT gray wolf embryos reconstructed with EPCs and **(B)** pericytes. **(C)** iSCNT plains bison embryos reconstructed with EPCs with three cavitating blastocysts and **(D)** one hatching blastocyst after seven days of culture **(E)** iSCNT plains zebra embryos reconstructed with EPCs with six compacting morulae by seven days of culture. **(F)** A transabdominal ultrasound image shows a sagittal image of one of six viable gray wolf fetuses at 23 days after ET.

*Bovids.* Blood-derived EPCs were compared directly to fibroblasts from the same female bison donor for iSCNT using in vitro matured cow oocytes. Of 86 bison EPC-derived iSCNT embryos, 69 embryos cleaved (80% cleavage rate) with five developing to blastocyst stage by day 7 (7% of cleaved embryos; Fig. 7C). In comparison, 67 of 87 fibroblast-derived iSCNT embryos cleaved (77% cleavage rate) with two progressing to blastocyst stage (3% of cleaved embryos; Fig. 7D). Genotyping confirmed that all bison iSCNT blastocysts were derived from bison donor cells.

*Equids.* iSCNT embryos were successfully generated using male plains zebra EPCs and fibroblasts as donor cells in enucleated domestic horse oocytes. Sixteen out of 51 iSCNT zebra embryos reconstructed with EPCs (Fig. 7E) developed to the morulae and blastocyst stages (32%) within 6 to 8 days of culture and were cryopreserved. In parallel, 13 out of 31 iSCNT zebra (42%) embryos produced with fibroblasts developed to the morulae and blastocyst stages prior to cryopreservation.

## V. Discussion

### Blood-derived cells offer practical advantages for conservation biobanking

This study demonstrates that endothelial progenitor cells (EPCs) and pericytes isolated from peripheral blood are a robust alternative to dermal fibroblasts for conservation biobanking and interspecific somatic cell nuclear transfer (iSCNT). Blood-derived cells exhibit 2-3 fold faster doubling rates (15-20 hours versus >35 hours for fibroblasts) and reduce the timeline for generating banked cell lines from 3-4 weeks to 1.5-2 weeks (Fig. 1, 2). Combined with genomic stability equivalent to tissue-derived cells (Fig. 5) and demonstrated viability for generating cloned embryos across three mammalian genera (Fig. 7), blood-derived cells offer practical advantages for accelerating genetic rescue efforts.

The collection of peripheral blood provides several operational advantages over skin biopsies for wildlife biobanking. Blood sampling during routine veterinary examinations, health assessments, or translocation procedures creates opportunistic collection windows without requiring additional animal restraint or separate sampling events.

Unlike dart biopsies, which yield minimal tissue often insufficient for both explant and digestion derivation methods, blood samples provide abundant starting material. The sterile, closed-system collection of blood in anticoagulant tubes contrasts with the microbial burden inherent to skin surfaces, which contributes to the higher contamination rates observed in explant cultures compared to blood-derived cultures.

The consistency of blood-derived cell doubling rates across phylogenetically diverse species (canids, bovids, equids) suggests broad taxonomic applicability and facilitates standardized biobanking workflows across diverse conservation programs. This contrasts with reported fibroblast derivation variability, such as elephant biopsies showing 0-80% success even within the same individual (Jansen van Vuuren et al., 2023). Whole blood samples are already routinely collected with anticoagulants for genomic studies and non-viably frozen in the field without cryoprotectants for DNA extraction. With advanced planning, blood collection protocols could preserve options for both immediate genomic analysis and future cell line establishment, maximizing the value of each sampling event. This dual-purpose approach is particularly relevant for cryptic, nocturnal, or otherwise difficult-to-sample species where opportunities for biological sample collection are rare.

### Molecular and functional characterization confirms distinct cell populations

Comprehensive proteomic profiling established that EPCs, pericytes, and fibroblasts are biologically distinct populations with lineage-specific molecular signatures (Fig. 6). Fibroblasts exhibited elevated ECM production markers (COL1A1, COL3A1, FAP), pericytes showed high abundance of adhesion and vascular remodeling proteins (ITGB5, ICAM1, MMP14, MCAM), and EPCs expressed endothelial-associated markers (ENG, CDH5, PDGFRA) along with canonical EPC markers (CD34, PECAM1, SELE, NOS3) detected exclusively in EPCs (Fig. 6, Table S1). The statistical significance of expression differences (Log₁₀(p-value) > 2.8) and consistency across biological replicates confirm that blood-derived cultures yield distinct cell types rather than heterogeneous mixtures.

For practical biobank characterization without mass spectrometry, morphological assessment combined with doubling rate measurements provides reliable first-pass identification (Fig. 2, 3). EPCs consistently display cobblestone morphology and ∼15-hour doubling times, while fibroblasts show spindle morphology with >35-hour doubling times. Pericytes exhibit intermediate characteristics with fibroblast-like morphology but faster doubling (∼20 hours). Functional assays further distinguish cell types within 24 hours: tubule formation on extracellular matrix occurs exclusively in EPC cultures, while differential uptake of fluorescently labeled LDL versus acetylated-LDL distinguishes EPCs from pericytes (Fig. 4). These species-agnostic methods should enable biobank curators to characterize cell lines from taxonomically diverse samples as collections expand to under-represented vertebrate lineages.

### Both EPCs and pericytes support viable iSCNT outcomes

Our iSCNT experiments demonstrate that blood-derived cells generate viable embryos with efficiency comparable to or exceeding traditional fibroblast donors. In gray wolves, both pericytes (19% implantation) and EPCs (11% implantation) contributed to viable fetuses, with the combined rate (15%) exceeding prior published wolf iSCNT rates using fibroblasts (M. K. Kim et al., 2007). The bison experiments provided direct within-individual comparison, as EPCs from the same female donor showed higher blastocyst formation (7%) than fibroblasts (3%) under identical conditions, suggesting that rapid proliferation and maintained progenitor characteristics may confer advantages for post-transfer embryonic development. These results overcome previously reported developmental blocks in bison iSCNT (González-Grajales et al., 2016). Plains zebra iSCNT embryos generated with EPCs (32% to morula/blastocyst) and fibroblasts (42%) further demonstrate cross-species applicability. Collectively, these results indicate that cell source (blood versus tissue) and specific blood-derived cell type do not compromise cloning outcomes.

### Current limitations and opportunities for optimization

The stochastic emergence of either EPC-dominant or pericyte-dominant cultures from blood samples presents both a challenge and an opportunity. In our study, replating the total buffy-coat fraction resulted in primary adherent colonies of both cell types adhering to traditional cell culture substrates and media optimized for EPC derivation and expansion. Both cell types formed primary colonies in a similar time window and could be dissociated, replated, and grown rapidly. While admixture of both cell types in cultures cannot be ruled out without single-cell analyses, the formation of cultures appeared stochastic with either EPCs or pericytes forming the predominant cell type in individual cultures. Selective enrichment may be achievable through substrate modification, growth factor supplementation, or differential adhesion protocols, enabling targeted derivation based on intended downstream applications. Both EPCs and pericytes participate in angiogenic signaling and repair and provide cell models for studying pathophysiological processes like atherosclerosis, making such optimization valuable for future studies beyond conservation applications.

Species-specific optimization of minimum blood volumes (currently 15 mL for canids, 25 mL for bovids and equids) requires further investigation across taxonomic groups. Body size, hematocrit, and circulating EPC/pericyte frequency likely influence volume requirements, and establishing taxon-specific guidelines would improve collection success rates while minimizing physiological stress to sampled individuals.

An important area for optimization is improving derivation success from cryopreserved field samples. All fresh tissue collections in this study were processed within 48 hours for cell line derivation. However, when animals under managed care could be safely restrained with advanced planning but transport to a laboratory within 48 hours was not feasible, we collected both blood and tissue to viably freeze in the field. We observed decreased success of derivation from cryopreserved samples (data not included), though successful preservation of the option for viable cell expansion at a later date remains a frequent real-world necessity. Developing robust field-to-lab workflows that maintain both genomic quality and cell viability would maximize the utility of limited sampling opportunities, particularly for endangered species where repeated sampling is not feasible.

### Integration with conservation strategies and existing biobanking infrastructure

The decision between blood and tissue sampling should be guided by species-specific considerations including restraint safety, veterinary access, and existing protocol success rates. For taxa where tissue biopsy success is inconsistent or anatomical considerations preclude safe sampling, blood collection offers clear advantages. The rapid expansion timeline has implications for time-sensitive interventions in species experiencing rapid population decline. Recent successes introducing cloned historically underrepresented founders into black-footed ferret and Przewalski’s horse populations (Novak et al., 2024; Novak, Ryder, et al., 2025) illustrate how biobanked cells restore genetic variation unavailable through conventional breeding. Blood-derived cells expand the pool of individuals who can be biobanked before unique genotypes are lost.

Integration with existing repositories such as the Frozen Zoo® is straightforward. Blood-derived cells can be processed using standard cryopreservation protocols and stored under identical conditions as tissue-derived lines. Unlike the technical demands to generate and biobank stem cells (Hutchinson et al., 2024), blood-derived cells can be processed using standard cryopreservation protocols and stored under identical conditions as tissue-derived lines. The addition of blood-derived samples would enhance representation of individuals for whom tissue biopsies were not collected, expanding the genetic diversity captured in these repositories. Our demonstrated success across phylogenetically distant mammalian orders suggests applicability to additional vertebrate taxa, potentially including threatened avian, reptilian, and amphibian species where current biobanking and conservation breeding efforts remain limited compared to the scale and rates of extinction (Browne et al., 2024).

However, the success of any biobanking-to-cloning effort depends critically on co-development of Assisted Reproductive Technologies (ART) using non-threatened, related model species whenever available and appropriate. Cloning of a new species can only happen when its basic reproductive physiology is understood and requires supporting the development of foundational ART to be successful. The iSCNT results reported here were only possible because of established ART protocols in domestic dogs, cattle, and horses, emphasizing that cloning advances depend on robust foundational reproductive biology research (Cowl et al., 2024; Hildebrandt & Holtze, 2024). Investing in studies and methods to collect viable gametes for artificial insemination, production of embryos by IVF, and timing of donors and recipients for successful embryo transfer can have profound positive impacts on preservation of current endangered populations (Bolton et al., 2022; Hutchinson et al., 2024).

Beyond technical advances in biobanking and ART, the selection of genetically valuable individuals for cloning should be performed in coordination with local and international community members invested in the propagation of endangered species and populations (Gray et al., 2023; Novak, Brand, et al., 2025). Importantly, supporting both international and indigenous groups advocating for *in situ* and *ex situ* conservation efforts of transnational, endangered animal populations and their habitats is needed in parallel to biobanking and long-term cloning efforts (Bolton et al., 2022; Hayah et al., 2025). These are complementary, not competitive, approaches. Engaging with motivated, multi-disciplinary parties may ultimately be more successful in achieving the shared goal to preserve more species from anthropogenic accelerated extinction.

By integrating blood-derived cell collection into routine management activities, conservation programs can opportunistically capture genetic diversity during standard veterinary procedures, expanding options for genetic rescue when traditional reproductive technologies cannot achieve sufficient genetic diversity. The minimal infrastructure requirements make this approach accessible to field programs with limited resources, broadening the pool of individuals and species represented in global biobanking efforts.

## VI. Data Availability

Proteomics data are available as Supplementary File 1.

## VII. Acknowledgements

We thank the UT Southwestern Proteomics Core Facility for their assistance in generating the mass spectrometry data, and KaryLogic Inc. for generating the canid karyotypes presented in this study. We also thank the organizations and owners who provided donations of samples collected from their privately owned animals.The domestic dog work described in this manuscript has been approved and conducted under the oversight of an independent Institutional Animal Care and Use Committee.

## VIII. Author Contribution and COI

JK, SB, TM, and BS conceived of the study. LK, AM, GG, RH, KMP, BS, and JK participated in data curation and formal analysis. BL provided financial support. JK AM, TM, FC, GLG, RH, JRH, LK-Y, LL, AB, SM, JBP. PP. LR, KR, KSr, KSt, DT, KW, RVS, SB performed experiments. TM, JH, SM, SB, MJ, SW, BS, and JK provided management/supervision and validated results. LK, AM, RH, KM-P, BS, and JK created visuals. JK, LK, and BS wrote the original draft of the manuscript, and all authors reviewed the final draft.

Authors affiliated with Colossal Biosciences may hold stock and/or stock options in the company.

## X. Supplemental Figures and Tables

**Figure S1.**
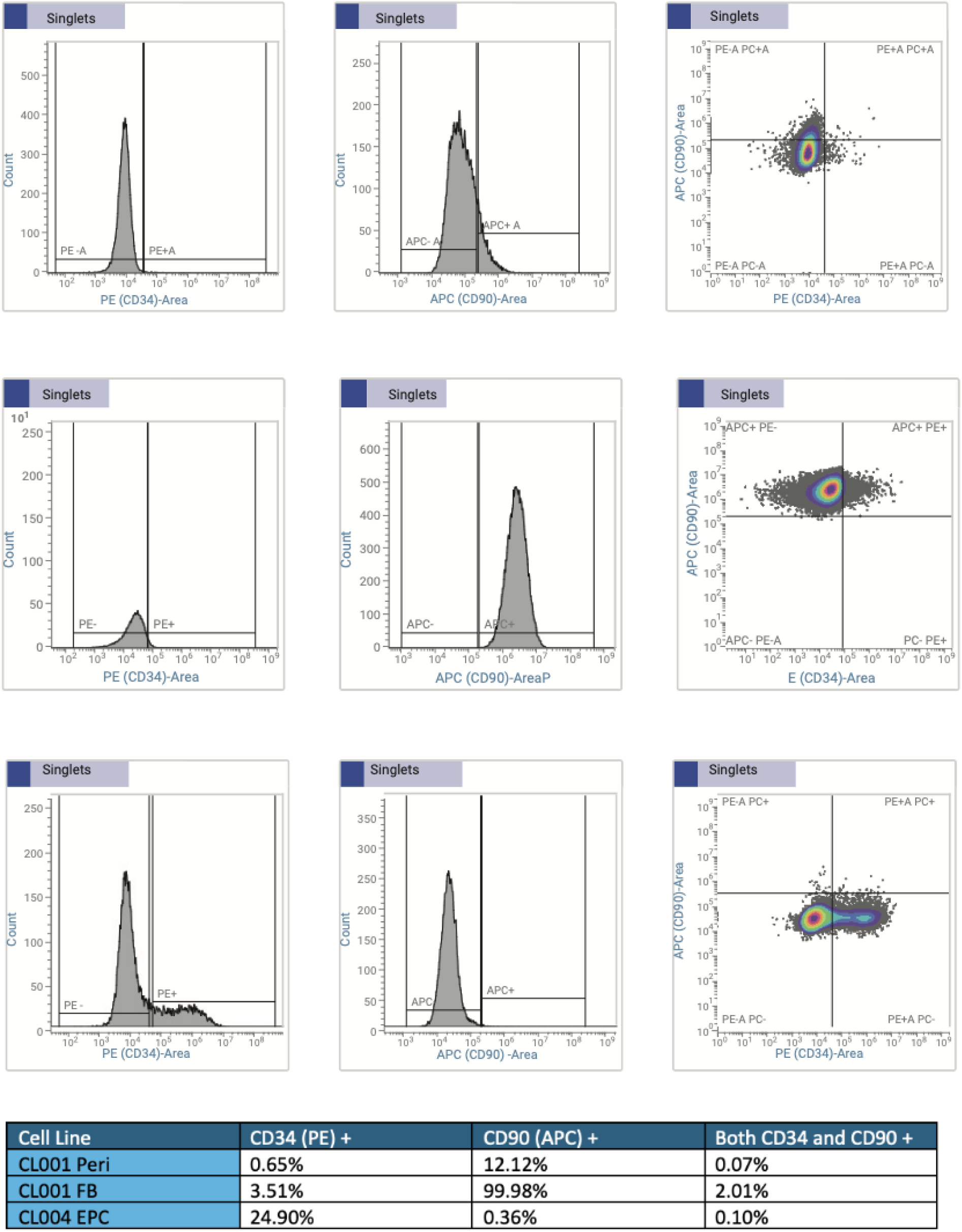
FACS analyses of CD34 and CD90 expression on gray wolf cell populations. A. Single parameter histograms and dual-parameter dot plots detected a 12% sub-population of CD34-, CD90+ canid pericytes. While, B. Canid fibroblasts were 100% CD90+ with a 3.5% sub-population co-expressing CD34 C. Canid EPCs had a discrete 25% sub-population that were CD34+ and all were CD90-. Forward and side scatter gating was used to collect data on singlets and exclude cell clusters, and thresholds were set using cells stained with isotype controls (not shown). Quadrant percentages for dot plots are summarized at the bottom.

**Figure S2.**
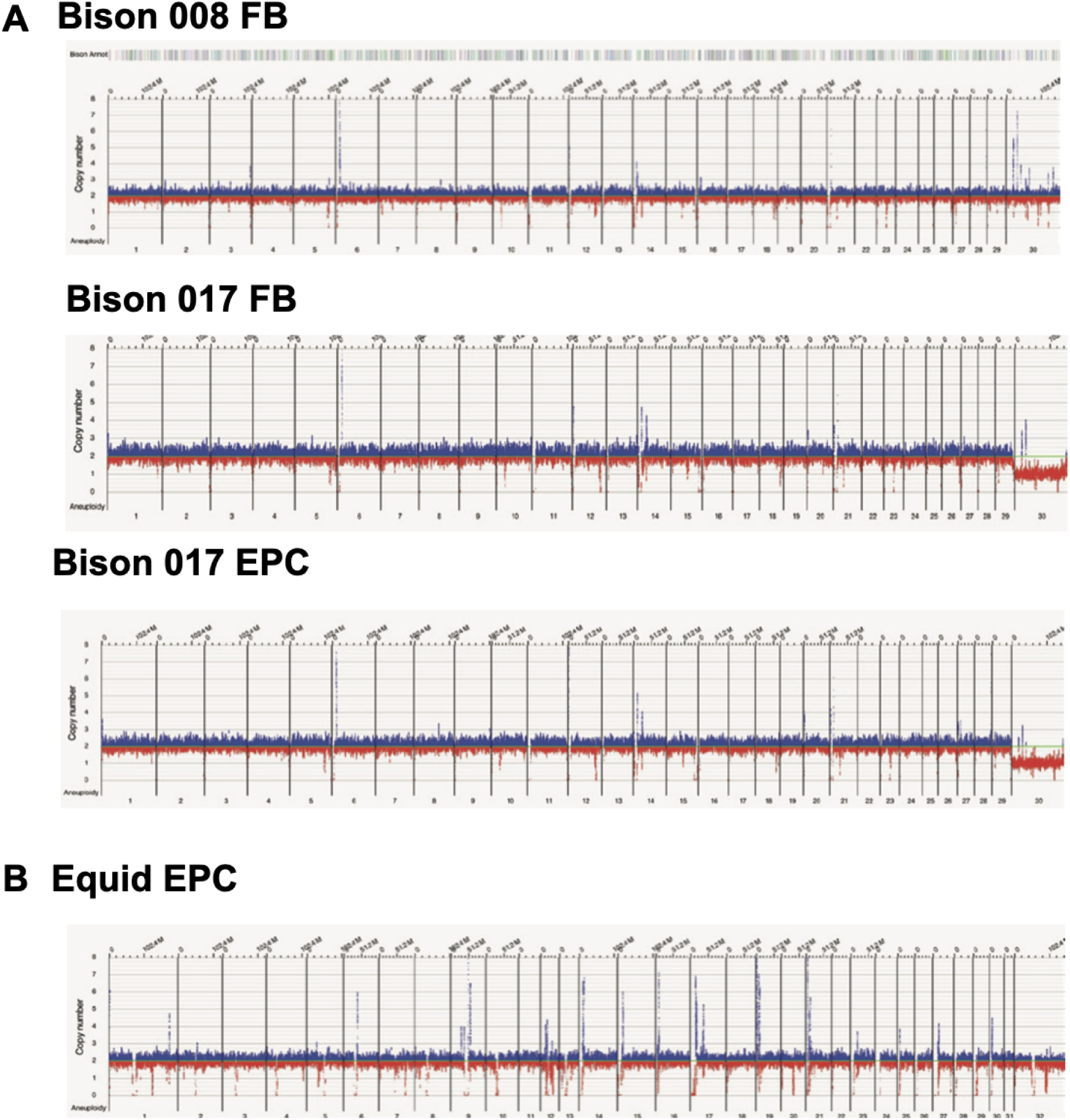
Optical Genome Mapping (OGM) plots summarizing chromosome copy numbers for **A.** bovid cell lines and **B.** a equid EPC line.

**Table S1.**
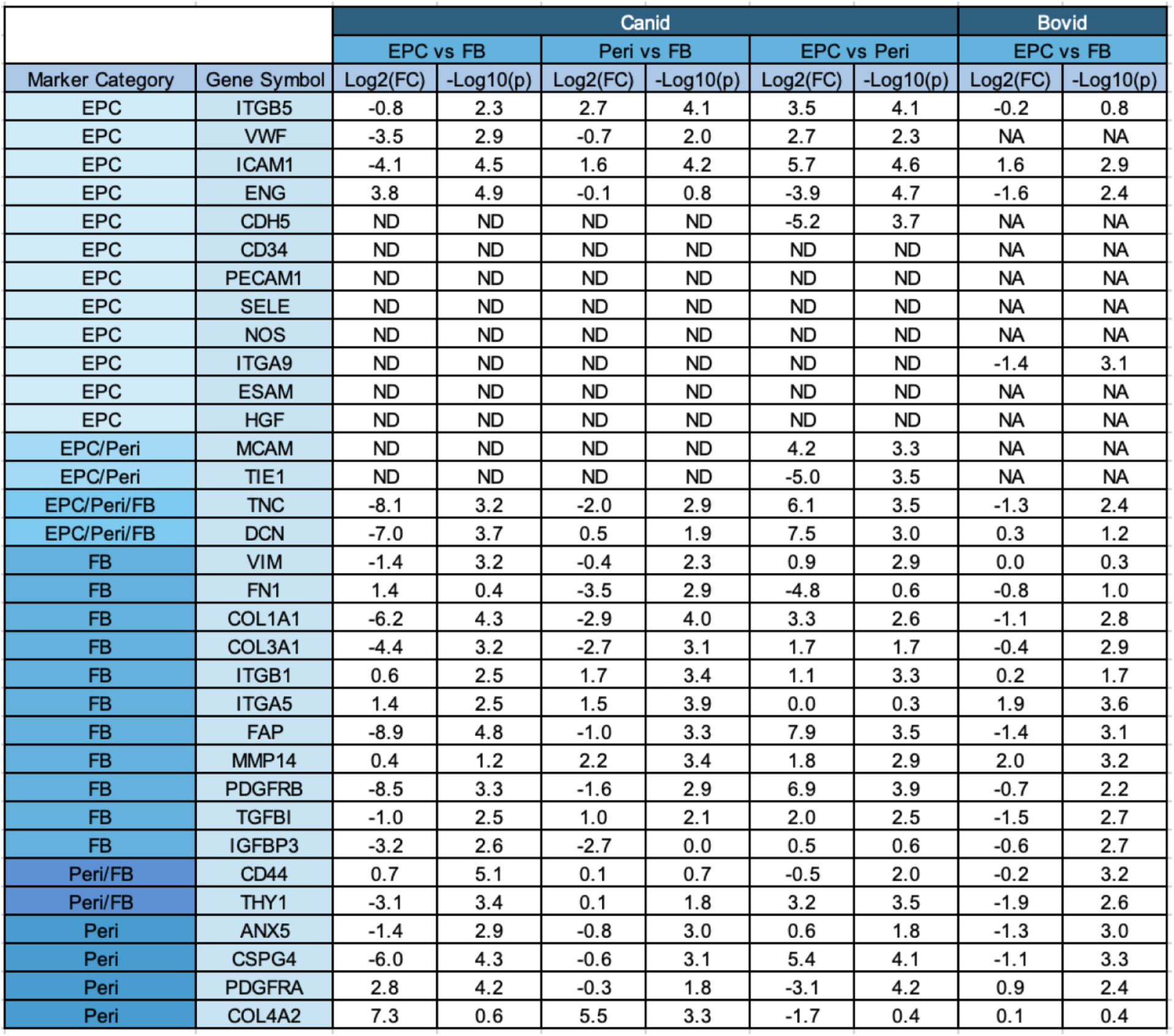
Log2(fold-change) and -Log10(p-value) for comparison between cell types for canonical marker-associated genes. A Log2(fold-change) value of 1 represents a 2-fold change in gene expression while a value of -1 indicates a 0.5-fold change. The -Log10(p-value) is a measure of statistical significance, where a value of 3 corresponds to a p-value of 0.001.

